# Investigating the temporal effects of ovarian failure on single muscle fibre contractility using a chemically-induced ovarian failure model in mice

**DOI:** 10.1101/2024.07.22.604625

**Authors:** Parastoo Mashouri, Jinan Saboune, W. Glen Pyle, Geoffrey A. Power

## Abstract

We investigated the effects of chemically-induced ovarian failure on single muscle fibre contractility of the soleus and extensor digitorum longus (EDL) muscles throughout ovarian failure, thereby mimicking the menopausal transition into late-stage menopause: [(D60;peri-menopause), (D120;onset of menopause), (D134;early-onset menopause), (D176;late-stage menopause)]. We used 4-vinylcyclohexene diepoxide (VCD) to induce ovarian failure in sexually-mature female mice. At D120 and D176, mice with VCD-induced ovarian failure produced higher force as compared with controls (p<0.05). On D134, however, VCD had lower force production compared with controls (p<0.05). The cross-sectional area of the soleus fibres from the VCD group was larger at D120 compared with controls (p<0.05), but not at any other time point (p>0.05). As well, at D120-D176, the proportion of Type II fibres relative to Type I increased for the soleus, but not the EDL. No differences in rate of force redevelopment (Ktr) was observed for the soleus (p>0.05), while calcium sensitivity increased by late-stage menopause (p<0.05). There were no differences in force, cross-sectional area, stiffness, Ktr, or calcium sensitivity between groups for the EDL (p>0.05). Muscle contractility across the peri-menopausal transition into late-stage menopause is both muscle and phase-dependent, emphasizing the complexity of changing hormones throughout the lifespan on muscle contractile function.

## Introduction

Perimenopause refers to the time when the female body begins the natural transition into menopause, which can take several years (El Khoudary et al., 2019). Menopause is defined by the permanent cessation of menstruation and is linked to a decrease in ovarian follicular activity, resulting in a reduction of 17β-estradiol production (Davis et al., 2015). This reduction in 17β-estradiol contributes to impaired static and dynamic muscle contractile function (Collins et al., 2019; Hubbard et al., 2023; Lowe et al., 2010; Sipilä & Poutamo, 2003; Tiidus, 2011; Tiidus et al., 2013). Most studies investigating the effects of reduced estrogens on muscle contractile function use the ovariectomized (OVX) model, whereby ovaries are surgically removed, leading to a sudden halt in ovarian hormone production which does not mimic the prolonged transitional phase of naturally occurring menopause (Hoyer et al., 2001; Mayer et al., 2004). Thus, in the present study, we induced gradual ovarian failure in mice to allow for the investigation of the perimenopausal transition on muscle contractile function (Brooks et al., 2016; Fernandes et al., 2019; Hubbard et al., 2023; Mashouri et al., 2024) into late-stage menopause (Hinks et al., 2024).

17β-estradiol-deficiency is generally associated with reductions in both absolute and specific (absolute force normalized to muscle size) force production in skeletal muscle (Collins et al., 2019; Lowe et al., 2010; Tiidus, 2011; Tiidus et al., 2013), but not always (Fillion et al., 2019; Greising, Baltgalvis, et al., 2011; Hinks et al., 2024; Mashouri et al., 2024). Using an OVX model of ovarian failure in adult rodents, reductions in absolute and specific force at the whole muscle and single fibre level by ∼10-30% for the soleus (SOL) and extensor digitorum longus (EDL) have been reported ∼1-3 months post-OVX (Greising, Baltgalvis, et al., 2011; Hubbard et al., 2023; Moran et al., 2007; Moran et al., 2006; Wattanapermpool & Reiser, 1999). However, others using similar models reported no difference in absolute or specific force for SOL and EDL whole muscles in ∼2 (Fillion et al., 2019) and ∼5-6-month-old rodents 2 weeks (Fillion et al., 2019; Greising, Baltgalvis, et al., 2011) or 2 months post-OVX (Cabelka et al., 2019; Cabelka et al., 2021; Hinks et al., 2024). Some groups have even reported absolute force values ∼20% higher for the EDL muscle at the joint level for 10-week-old rodents 7 weeks post-OVX compared to age-matched controls (Suzuki & Yamamuro, 1985). Using a VCD-induced model of ovarian failure, the absolute and specific force of the SOL whole muscle and plantar flexors in young mice was not different (Greising, Carey, et al., 2011; Hinks et al., 2024). However, in our previous investigations using the VCD model, we found that single fibre absolute force in young, sexually mature mice at the onset of menopause was higher in the VCD group in the SOL but not the EDL (Mashouri et al., 2024). We related these muscle-dependent findings to a shift in fibre type from slow to fast in the SOL muscle (Mashouri et al., 2024; Moran et al., 2007). Cross-bridge-based mechanisms are suggested to contribute to impaired muscle force production in both OVX and VCD-induced models of ovarian failure (Greising, Carey, et al., 2011; Mashouri et al., 2024). While some results indicate that reductions in circulating 17β-estradiol levels are associated with reductions in strongly bound cross-bridge states by ∼10-15% (Moran et al., 2007; Moran et al., 2006; Peyton et al., 2022), resulting in less force per myosin-actin interaction, others have shown no change in the proportion of attached cross-bridges (Mashouri et al., 2024). Moreover, myofibrillar calcium (Ca^2+^) sensitivity does not seem to contribute to force loss in the OVX model for the SOL muscle (Wattanapermpool & Reiser, 1999), nor the VCD model for both the EDL and SOL muscles at the onset of menopause (Mashouri et al., 2024). These divergent findings warrant further investigation as the different models, rodent ages, and time course used may explain the between-study variability observed in the literature.

Most studies investigating post-menopausal changes in skeletal muscle contractility have used OVX rodent models. Although OVX produces a 17β-estradiol-deficient state like that experienced by post-menopausal women, it fails to simulate the gradual onset, extended transitional period, and preservation of ovarian tissue typical of menopause in most women. As a result, the failure to reproduce these physiological features of menopause in animal experiments restricts the generalizability of this research to the human population (Fernandes et al., 2019). Thus, we investigated the effects of a gradual transition from perimenopause into late menopause phases using a VCD-induced ovarian failure model in mice to determine time-course effects of menopause on skeletal muscle contractility. While our previous findings using the VCD model indicated an increase in absolute force for the SOL muscle at the onset of ovarian failure, we believe as ovarian failure progresses the prolonged 17β-estradiol-deficient state will have divergent results. We hypothesize that as ovarian failure progresses, i) absolute force, specific force, instantaneous stiffness and rate of force redevelopment (Ktr) will be lower over time from the perimenopausal transition into late-stage menopause; ii) the SOL muscle will have a greater response to 17β-estradiol deficiency compared to the EDL muscle; and iii) Ca^2+^ sensitivity will not differ between groups but will vary in a muscle-dependent manner.

## Methods

### Animals

Sexually mature CD1 female mice (n=30; n=15 VCD; n=15 control) aged 78 105 days were obtained from Charles River Laboratories (St. Constant, QC), and housed on a 12 h light/dark cycle. Food and water were provided ad libitum. All procedures conducted were approved (AUP: 4714) and performed in accordance with the guidelines set by the Animal Care and Use Committee of the University of Guelph and the Canadian Council on Animal Care.

### 4□Vinylcyclohexene diepoxide (VCD) mouse model of menopause

Mice were weighed and given daily intraperitoneal injections (160mg/kg) of VCD (MilliporeSigma, Oakville, ON) for 15 consecutive days (Hinks et al., 2024; Hubbard et al., 2023; Mashouri et al., 2024; Perez et al., 2013). VCD accelerates follicular atresia by inducing the selective loss of primary and primordial follicles in the ovary but leaves intact the rest of the ovary for residual hormone production. VCD mediates its ovarian effects by inhibiting autophosphorylation of membrane receptor c□kit. Vaginal cytology was used to confirm lack of estrus cycles indicating ovarian failure (Caligioni, 2009; Cora et al., 2015). Mice were considered acyclic after 10 consecutive days of persistent diestrus which was achieved by day 120 representing the end of perimenopause and the start of menopause (Fernandes et al., 2019). Muscle samples from all VCD mice were taken at days 60, 120, 134 and 176 following the start of VCD injections, representing the different stages of menopause: perimenopause; onset of menopause; early onset-menopause; and late onset-menopause, respectively.

### Tissue Preparation

The SOL and EDL muscles were harvested following sacrifice and placed in a chilled dissecting solution, chemically permeabilized, and stored as reported previously (Hubbard et al., 2023; Mashouri et al., 2024).

### Solutions

The dissecting solution was composed of the following (in mM): K proprionate (250), Imidazole (40), EGTA (10), MgCl_2_·6H_2_O (4), Na_2_H_2_ATP (2), H_2_O. The storage solution was composed of the following (in mM): K proprionate (250), Imidazole (40), EGTA (10), MgCl_2_·6H_2_O (4), Na_2_H_2_ATP (2), glycerol (50% of total volume after transfer to 50:50 dissecting:glycerol solution), as well as leupeptin (Sigma) protease inhibitor. The skinning solution with Brij 58 was composed of the following (in mM): K proprionate (250), Imidazole (40), EGTA (10), MgCl_2_·6H_2_O (4), 1 g of Brij 58 (0.5% w/v). The relaxing solution was composed of the following (in mM): Imidazole (59.4), K.MSA (86), Ca(MSA)2 (0.13), Mg(MSA)_2_ (10.8), K3EGTA (5.5), KH_2_PO_4_ (1), H_2_O, Leupeptin (0.05), Na_2_ATP (5.1), as well as leupeptin (Sigma) protease inhibitor. The pre□activating solution was composed of the following (in mM): KPr (185), MOPS (20), Mg(CH_3_COOH)_2_ (2.5), ATP (2.5). The activating solutions were composed of the same ingredients in various amounts depending on how much Ca^2+^ was needed in each solution. The activating solution was composed of the following (in mM): Ca^2+^, Mg, EGTA (15), MOPS (80), ATP (5), CP (15), K (43.27), Na, and H_2_O. Ca^2+^, Mg, and Na concentrations were specific to the pCa level (bolded number): (i) **4.5**: Ca^2+^ (14.87), Mg (6.93), Na (13.23); (ii) **5.5**: Ca^2+^ (12.83), Mg (6.97), Na (13.4); (iii) **5.7**: Ca^2+^ (11.83), Mg (7), Na (13.4); (iv) **6.2**: Ca^2+^ (8.1), Mg (7.07), Na (13.4); (v) **6.4**: Ca^2+^ (6.4), Mg (7.07), Na (13.4); (iv) **6.6**: Ca^2+^ (4.8), Mg (7.1), Na (13.4); (iiv) **7.0**: Ca^2+^ (2.37), Mg (7.17), Na (13.4).

All solutions were adjusted to a pH of 7.0 with the appropriate acid (HCl) or base (KOH). The composition of solutions was determined by calculating the equilibrium concentration of ligands and ions based on published affinity constants (Fabiato and Fabiato, 1979). 250 units/ml of creatine phosphokinase was used in each activating solution. The homogenization buffer was composed of the following: 61 mM tris (pH 6.8), 11% (*v/v*) glycerol, 2.78% (*w*/*v*) SDS, 5% 2-β-mercaptoethanol, and 0.02% (*w/v*) bromophenol blue. The SDS-PAGE solutions were composed of the following: The 7% separating gel consisted of 2 M Tris HCL (pH 8.6), 50% glycerol, 10% sodium dodecyl sulfate (SDS), and 40% (*w/v*) acrylamide and N,N′-methylenebis-acrylamide with a monomer to crosslinker ratio of 37.5:1. The stacking gel consisted of 500 mM Tris HCL (pH 6.7), 10% SDS, and 40% (w/v) acrylamide and N,N′-methylenebis-acrylamide with a monomer to crosslinker ratio of 37.5:1.

### Mechanical testing and force measurements

Single fibres were transferred into a temperature controlled chamber (15°C) and tied to pins mounted between a force transducer (model 403A; Aurora Scientific) and a length controller (model 322C; Aurora Scientific). Average sarcomere length (SL) was measured using a high speed camera (Aurora Scientific). Fibre length (L_0_) was recorded, and fibre diameter was measured at three different points along the fibre using a reticule on the microscope in relaxing solution to calculate cross-sectional area (CSA) assuming circularity. First, a ‘fitness’ contraction was performed at 2.5 μm in pCa 4.5, after which SL was re-measured and, if necessary, re adjusted to optimal rodent hindlimb SL, 2.5 μm. To initiate activation, the fibres were transferred to a pre activating solution (reduced Ca^2+^ buffering capacity with ATP) for 20 s, then to an activating solution (varying levels of Ca^2+^ and high ATP) for 30 s (Mashouri et al., 2024). The highest force reached during the 30 s window was taken as optimal force (P_o_) at each pCa level.

To determine the force-pCa relationship, 7 pCa levels (7.0-4.5) were used in a random order. Data were fit using a modified Hill equation:

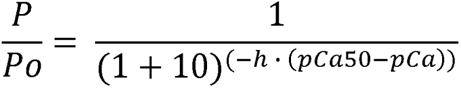

The pCa value at which 50% of maximal force was elicited (pCa_50_) was used to determine differences in Ca^2+^ sensitivity.

Ktr was assessed using a slack re-stretch method (Brenner & Eisenberg, 1986; Power et al., 2016). By rapidly shortening the fibre by 15% of L_o_ with a ramp of 10 L_o_/s and then a rapid (500 L_o_/s) re-stretch back to L_o_, we force all cross-bridges to break, and then the re-stretch allows further dissociation of any remaining cross-bridges. The force redevelopment is related directly to the reattachment of myosin to actin and a redistribution of cross bridges from pre-power stroke into force-generating states. A mono-exponential equation, y=a(1-e^-kt^)+b, was fit to the redevelopment curve to determine Ktr.

To assess the proportion of attached cross-bridges, instantaneous stiffness tests were performed by inducing a rapid (500 L_o_/s) stretch of 0.3% of L_o_ and dividing the change in force during the stretch by the length step. Instantaneous stiffness was calculated as the difference between peak force during the stretch and the average force over the 500 ms immediately prior to the stretch, divided by the change in fiber length induced by the stretch. Absolute force was reported as the average force over 500 ms during the steady-state phase. All force measures were adjusted for resting passive tension by subtracting the average baseline value for the first 50 ms of the trial in the relaxing solution, thus for all contractions, active force is reported. If at low [Ca^2+^] a negative value was recorded owing to any force drift of the transducer, the value was replaced with ‘0’. All force values were normalized to CSA for calculations of specific force.

If final maximum force decreased by >10% from the maximal force produced at the beginning of the experiment, or if the striation pattern of the muscle fibres became unclear such that measurements of sarcomere length were not possible, the experiment was discarded. Any fibre that slipped or ripped before the completion of the tests was removed from the testing apparatus and excluded. A total of N=163 fibres from the EDL and N=174 fibres from the SOL were included in the analysis.

### Fibre Typing

Following mechanical testing, fibre typing was performed as described previously (Hubbard et al., 2022; Mashouri et al., 2021; Mazara et al., 2021). Gels were run at a constant voltage of 50V for approximately 40h at 4°C. Fibres containing MHC I and MHC II were classified as Type I (TI) and Type II (TII) and were determined by comparing them to a standard protein ladder (Bio-Rad Protein Plus Standard 10-250 kD) with known molecular weights.

### Statistical Analysis

2-way ANOVAs (IBM SPSS Statistics, V29.0.1.0) were used to assess differences between groups (VCD, control) and across timepoints (D60, D120, D134, D176) for absolute force, specific force, Ktr, instantaneous stiffness, fibre CSA, fibre types (within SOL), pCa_50_ values. An alpha level of 0.05 was used. Data in the text are presented as a % difference of means. Figures are presented as Mean ± SEM.

## Results

### Soleus muscle

#### Absolute force

For absolute force, there was an interaction of group × timepoint (p<0.001), and a main effect of timepoint (p<0.001), with no main effect of group (p>0.05). Fibres in the VCD groups at timepoints D120 and D176 produced ∼25%, and ∼29% more force, respectively, and at timepoint D134 produced 61% less force than controls. Fibres at timepoint D120 produced ∼60% and ∼74% greater force compared to timepoints D60 and D176, respectively, and timepoint D134 produced ∼41% and ∼68% more force compared to timepoints D60 and D176, respectively (Figure 1A).

**Figure 1.**
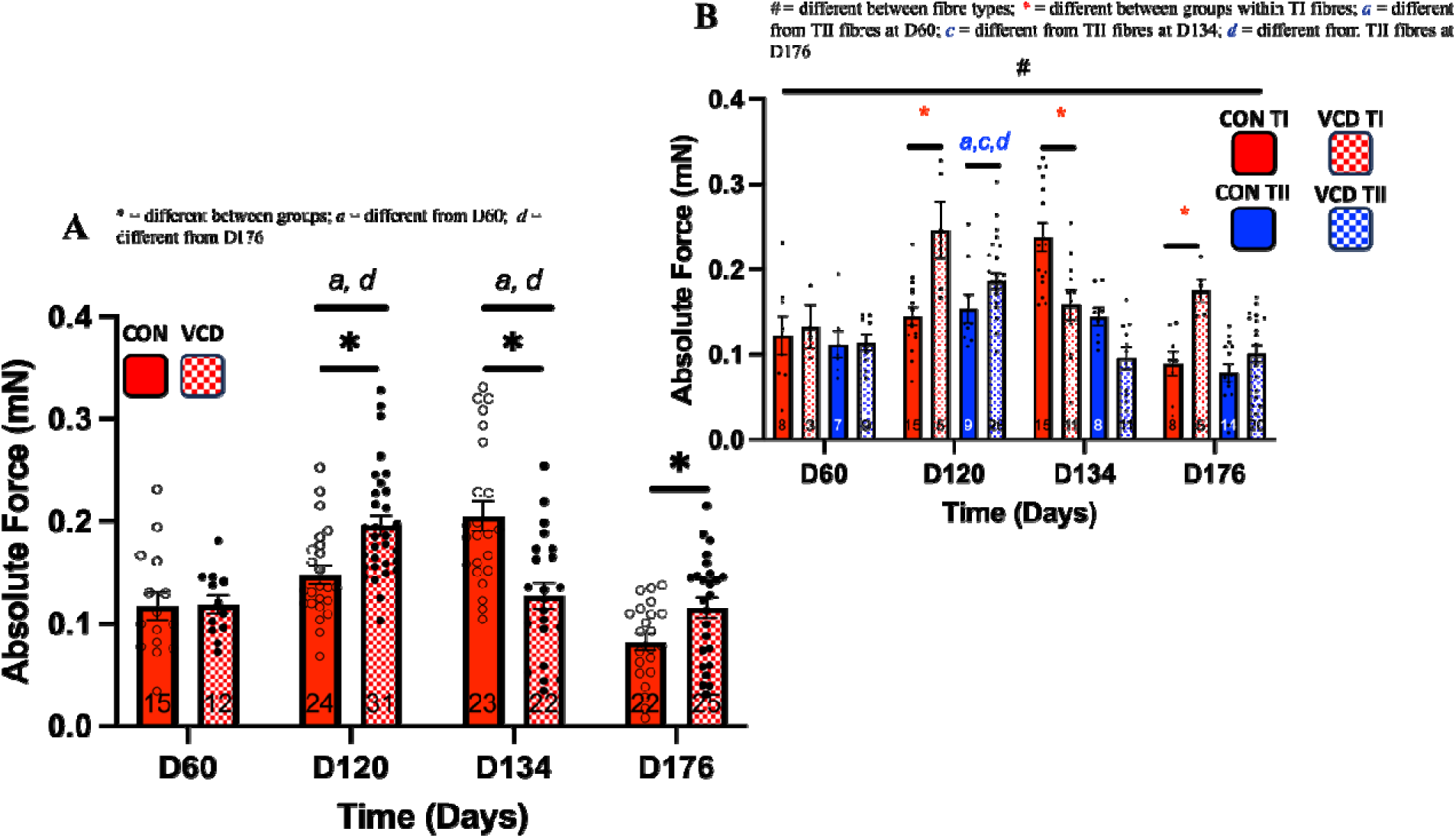
SOL Absolute force: A - Fibres from the VCD groups at D120 and D176 produced more force but at D134 produced less force than their controls. Across groups, fibres at D120 produced greater force compared to D60 and D176, and at D134 produced more force compared to D60 and D176. B – SOL muscle separated by fibre type. TI fibres from the VCD groups at D120 and D176 produced more force and at D134 produced less force than their controls, respectively. Additionally, TII fibres produced more force at D120 compared to all other timepoints. Lastly, at TI fibres produced more force than TII fibres across all timepoints.

When taking fibre type into consideration, there were no interactions of group × fibre type (p>0.05), timepoint × fibre type (p>0.05) or main effect of group (p>0.05). However, there was an interaction of group × timepoint × fibre type (p<0.05) such that TI fibres from the VCD groups at timepoints D120 and D176 produced ∼71% and ∼97% more force and at timepoint D134 produced ∼38% less force than their controls, respectively. There was also an interaction of group × timepoint (p<0.001), such that fibres in the VCD groups at timepoints D120 and D176 produced ∼45% and ∼64% more force and at timepoint D134 produced ∼34% less force than their controls, respectively. Additionally, fibres in the control group at timepoint D120 produced ∼77% more force than at timepoint D176, and fibres in the control group at timepoint D134 produced ∼28-127% more force compared to all other timepoints. Interestingly, fibres in the VCD group at timepoint D120 produced ∼57-76% more force than all other timepoints. There was also a main effect of timepoint (p<0.001), such that fibres at timepoint D120 produced ∼53% and ∼65% more force and fibres at timepoint D134 produced ∼33% and ∼43% more force compared to timepoints D60 and D176, respectively. Lastly, there was a main effect of fibre type (p<0.001), such that across all timepoints, TI fibres produced 40% more force compared to TII fibres (Figure 1B).

#### Cross-sectional area

For CSA, there was an interaction of group × timepoint (p<0.05) such that at timepoint D120, the VCD group had a ∼50% larger CSA compared with the CON group, and the CON group at timepoint D134 was ∼50% larger than at timepoints D60 and D176. There was also a main effect of timepoint (p<0.01) such that timepoint D134 had ∼50% larger CSA compared to timepoint D176 (Figure 2A).

**Figure 2.**
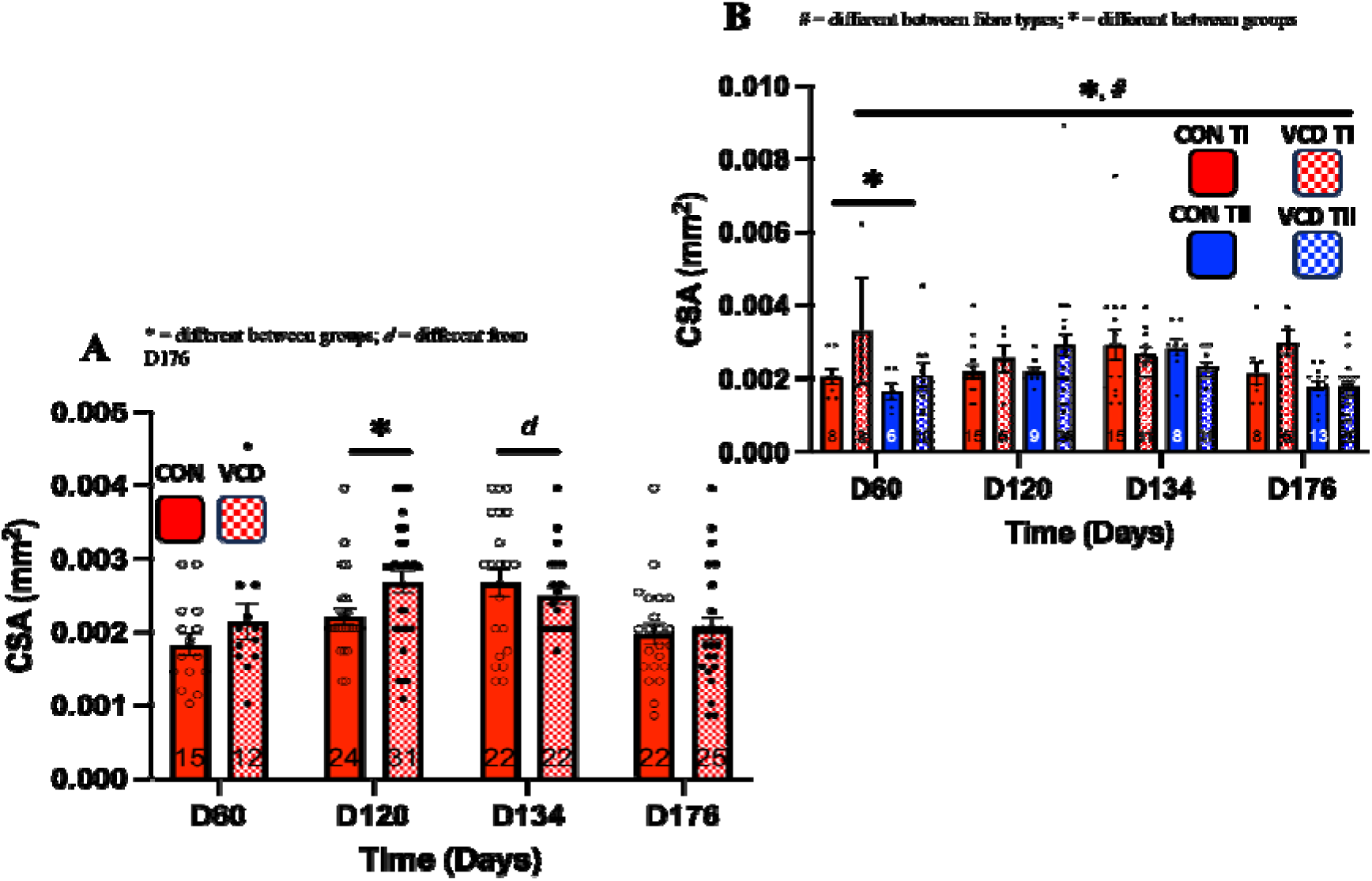
SOL CSA: A - At timepoint D120, the VCD group had larger CSA compared to the CON group. D134 had larger CSA compared to timepoint D176. B - SOL muscle separated by fibre type. The VCD group at timepoint D60 had larger CSA compared to controls. Additionally, the VCD group had larger CSA than controls. Lastly, TI fibres had larger CSA than TII fibres across all timepoints.

When taking fibre type into consideration, there was an interaction of group × timepoint (p<0.05), such that the VCD group at timepoint D60 had ∼50% larger CSA compared to controls and fibres in the CON group at timepoint D134 had ∼50% larger CSA than at timepoints D60 and D176. There was also a main effect of group (p<0.05) such that the VCD groups had a ∼50% larger CSA than the CON group. Lastly, there was a main effect of fibre type (p<0.05) such that TI fibres had ∼50% larger CSA than TII fibres across all timepoints (Figure 2B).

#### Specific force

For specific force, there was an interaction of group × timepoint (p<0.001), and an effect of timepoint (p<0.001) with no main effect of group (p>0.05). At timepoint D176, fibres produced ∼29% and ∼8% less specific force compared to timepoints D120 and D134, respectively. Additionally, fibres from the VCD group produced ∼37% less and ∼39% more force compared to controls at timepoints D134 and D176, respectively (Figure 3A).

**Figure 3.**
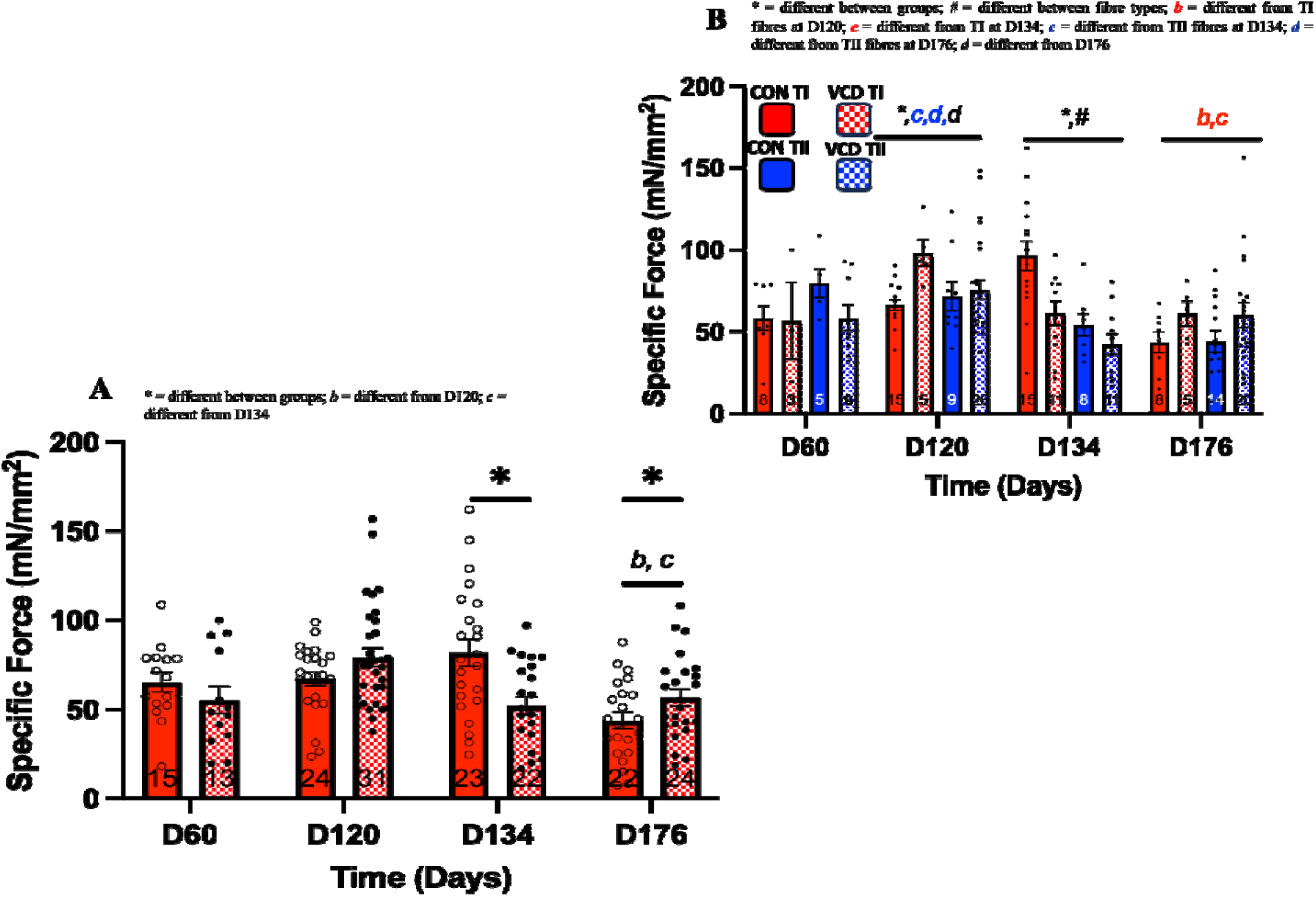
SOL Specific force: A - At timepoint D176, fibres produced less specific force across groups compared to timepoints D120 and D134. Additionally, fibres in the VCD group produced less and more force compared to controls at timepoints D134 and D176, respectively. B – SOL muscle separated by fibre type. At timepoint D120, the VCD group produced more specific force than the CON group, and at timepoint D134, the CON group produced more specific force than the VCD group. At timepoint D176, TI fibres produced less force compared to timepoints D120 and D134. Additionally, at timepoint D120, TII fibres produced more force than D134 and D176. At timepoint D134, TI fibres produced more force compared to TII fibres. Lastly, at timepoint D120 produced more specific force compared to timepoint D176.

When taking fibre type into consideration, there was an interaction of group × timepoint (p<0.001), such that at timepoint D120 the VCD group produced ∼26% more specific force than the CON group, and at timepoint D134, the CON group produced ∼45% more specific force than the VCD group. There was also an interaction of timepoint × fibre type (p<0.05) such that at timepoint D176, TI fibres produced ∼36% and ∼34% less force compared to timepoints D120 and D134, respectively. Additionally, at timepoint D120, TII fibres produced ∼52% and ∼41% more force than at timepoints D134 and D176, respectively. At timepoint D134, TI fibres produced ∼64% more force compared to TII fibres. Lastly, there was a main effect of timepoint (p<0.001) such that at timepoint D120, there was ∼49% more specific force compared to timepoint D176 (Figure 3B).

#### Instantaneous stiffness

For stiffness, there was an interaction of group × timepoint (p<0.01) and an effect of timepoint (p<0.01), with no main effect of group. At timepoint D120, there was 40% greater stiffness compared to at timepoint D176, indicating a higher proportion of attached cross-bridges. Additionally, at timepoint D134, fibres from the VCD group had ∼27% lower stiffness compared to controls, and at timepoint D176, this relationship was reversed, with the VCD group having ∼56% higher stiffness compared to controls (Figure 4A).

**Figure 4.**
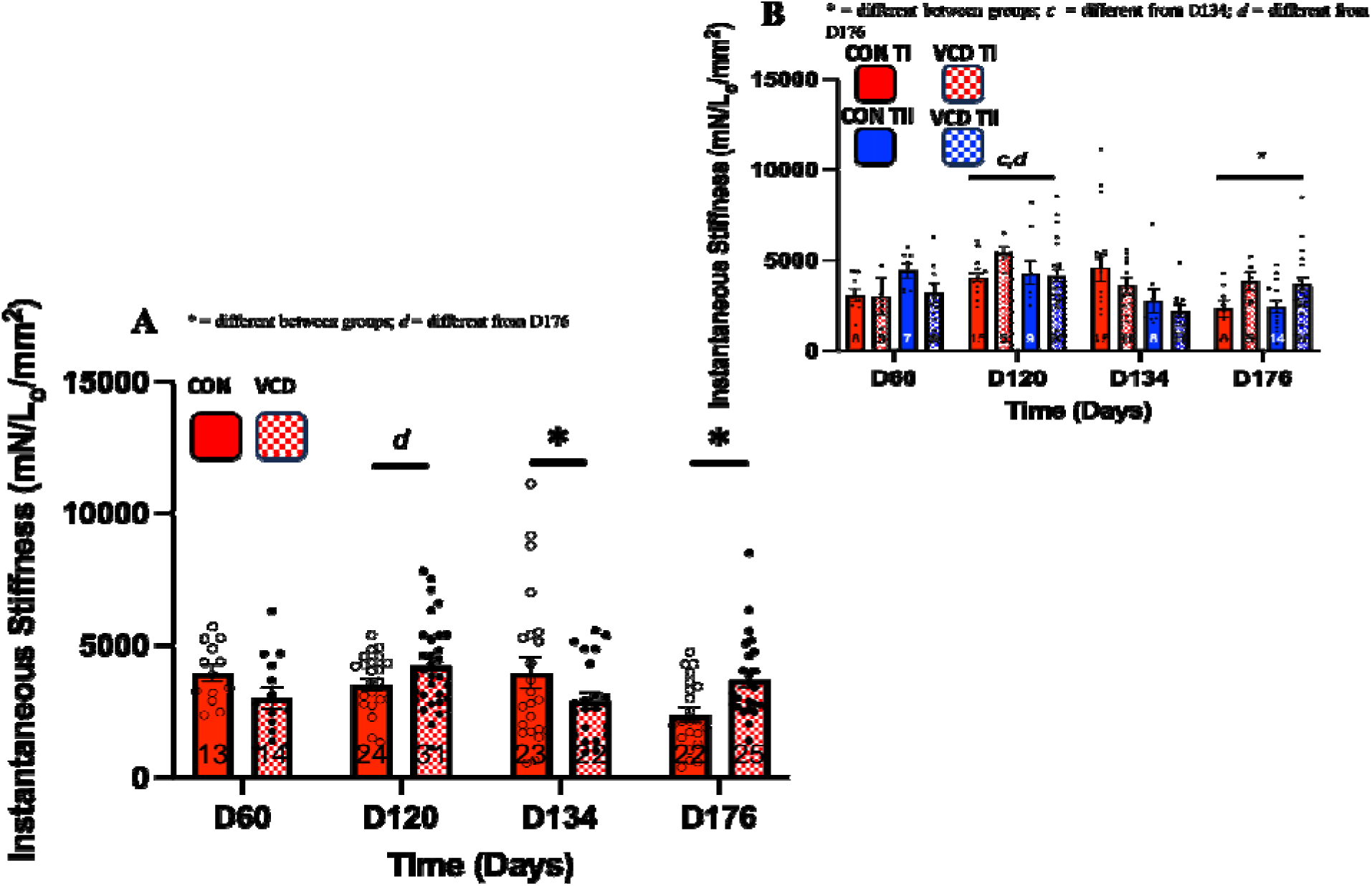
SOL Instantaneous stiffness: A – At timepoint D120, there was greater stiffness across groups compared to timepoint D176, indicating a higher proportion of attached cross-bridges. Additionally, at timepoint D134, fibres from the VCD group had lower stiffness compared to controls, and by timepoint D176, this relationship was reversed, with the VCD group having higher stiffness compared to controls. B – SOL muscle separated by fibre type. At timepoint D176, fibres from the VCD group had greater stiffness compared to the CON group. Lastly, fibres at timepoint D120 had higher stiffness compared to timepoints D134 and D176.

When taking fibre type into consideration, there was an interaction of group × timepoint (p<0.05) such that at timepoint D176, fibres from the VCD group had ∼59% higher stiffness compared to controls. Additionally, fibres from the CON group at timepoint D176 had ∼43% less stiffness than at timepoint D120 and fibres from the VCD group at timepoint D134 had ∼12% less stiffness than at timepoint D120. Lastly, there was a main effect of timepoint (p<0.01) such that fibres at timepoint D120 had ∼36% and ∼46% higher stiffness compared to timepoints D134 and D176 (Figure 4B).

#### Ktr

For Ktr, there was no interaction of group × timepoint or effect of group (p>0.05). However, there was a main effect of timepoint (p<0.001) such that timepoint D134 had ∼40% and ∼46% slower rates of force redevelopment compared to timepoints D60 and D176 (Figure 5A).

**Figure 5.**
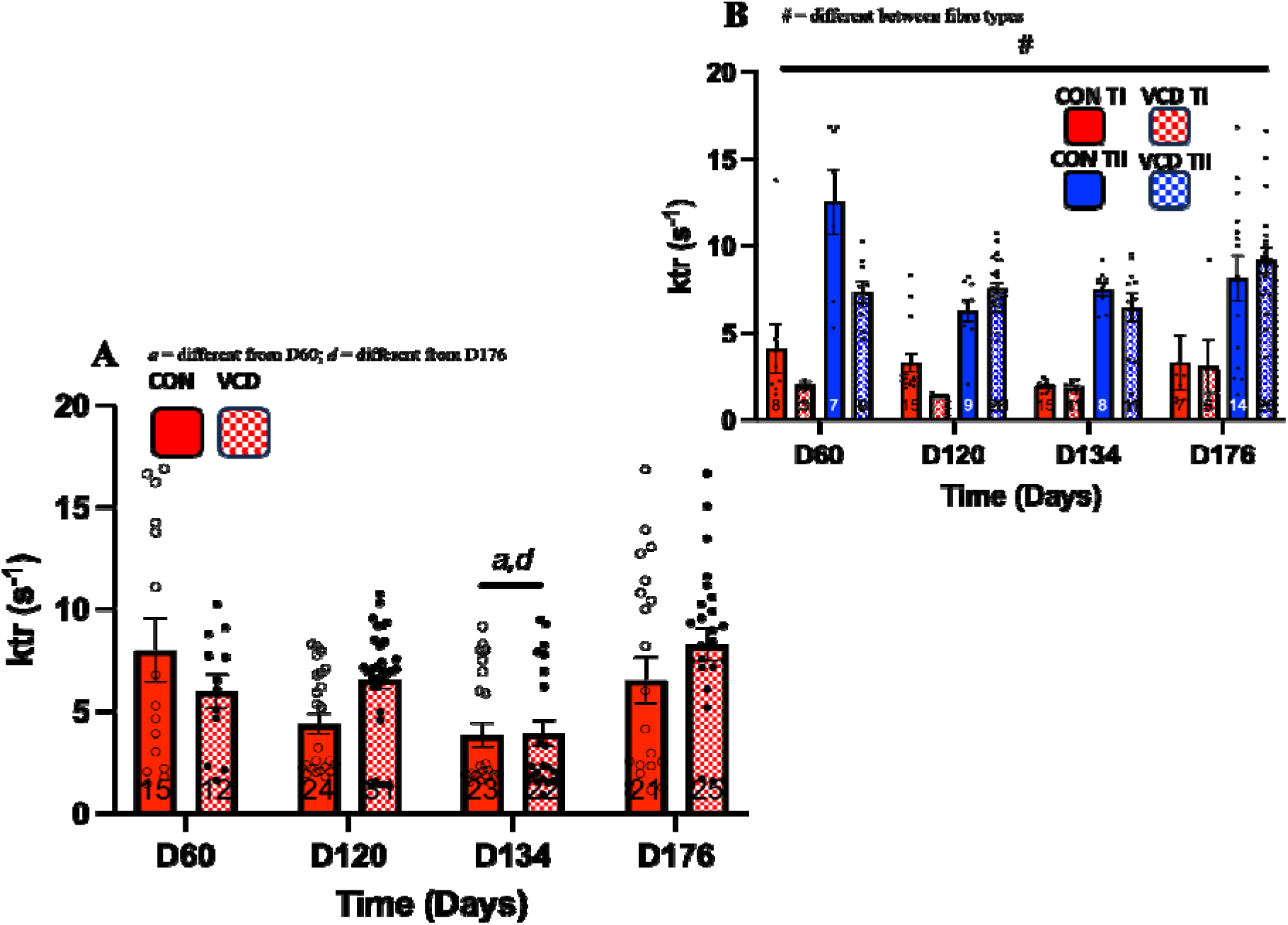
SOL Ktr: A – At timepoint D134, fibres across groups had a slower rate of force redevelopment compared to timepoints D60 and D176. B - SOL muscle separated by fibre type. Across groups, TII fibres produced faster Ktr than TI fibres.

When taking fibre type into consideration, there were no interactions (p>0.05). However, there was a main effect of timepoint (p<0.05) and a main effect of fibre type (p<0.001), such that TII fibres produced ∼200% faster Ktr than TI fibres (Figure 5B).

#### pCa_50_

For pCa_50_, there was an interaction of group × timepoint (p<0.001) such that at timepoints D60, D134 and D176, the VCD groups had ∼10%, ∼6% and ∼10% lower pCa_50_ values compared to controls, respectively, suggesting higher calcium sensitivity. There was a main effect of group (p<0.001) such that fibres from the VCD groups had a ∼6% lower pCa_50_ value compared to controls. There was also a main effect of timepoint (p<0.001) such that at timepoint D120, there were ∼4% and ∼3% lower pCa_50_ values compared to D60 and D176, and D134 had ∼5 and ∼4% lower pCa_50_ values compared to D60 and D176 (Figure 6).

**Figure 6.**
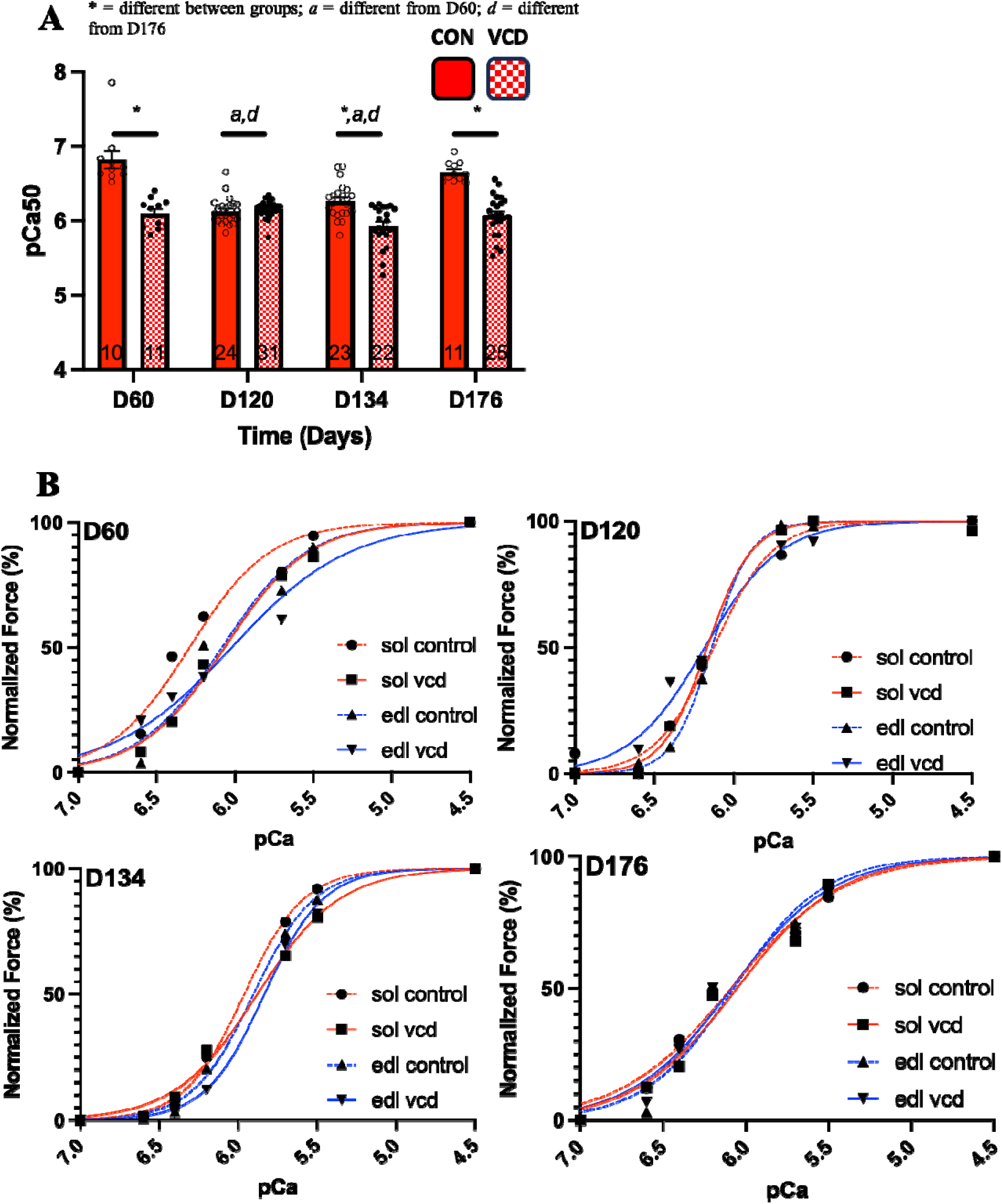
Calcium sensitivity: A - For pCa_50_, at timepoints D60, D134 and D176, fibres from the CON groups had greater calcium sensitivity compared to the VCD groups. At timepoint D120 fibres had a shift to higher calcium concentrations compared to timepoints D60 and D176. B – The force-pCa curves of both SOL and EDL at each timepoint are illustrated, showing the shifts in the midpoint of the curves, with fibres from the VCD group at timepoint D120 having lower calcium sensitivity compared to the CON group. All other timepoints had higher calcium sensitivity for the VCD group compared to the CON group.

**Figure 7.**
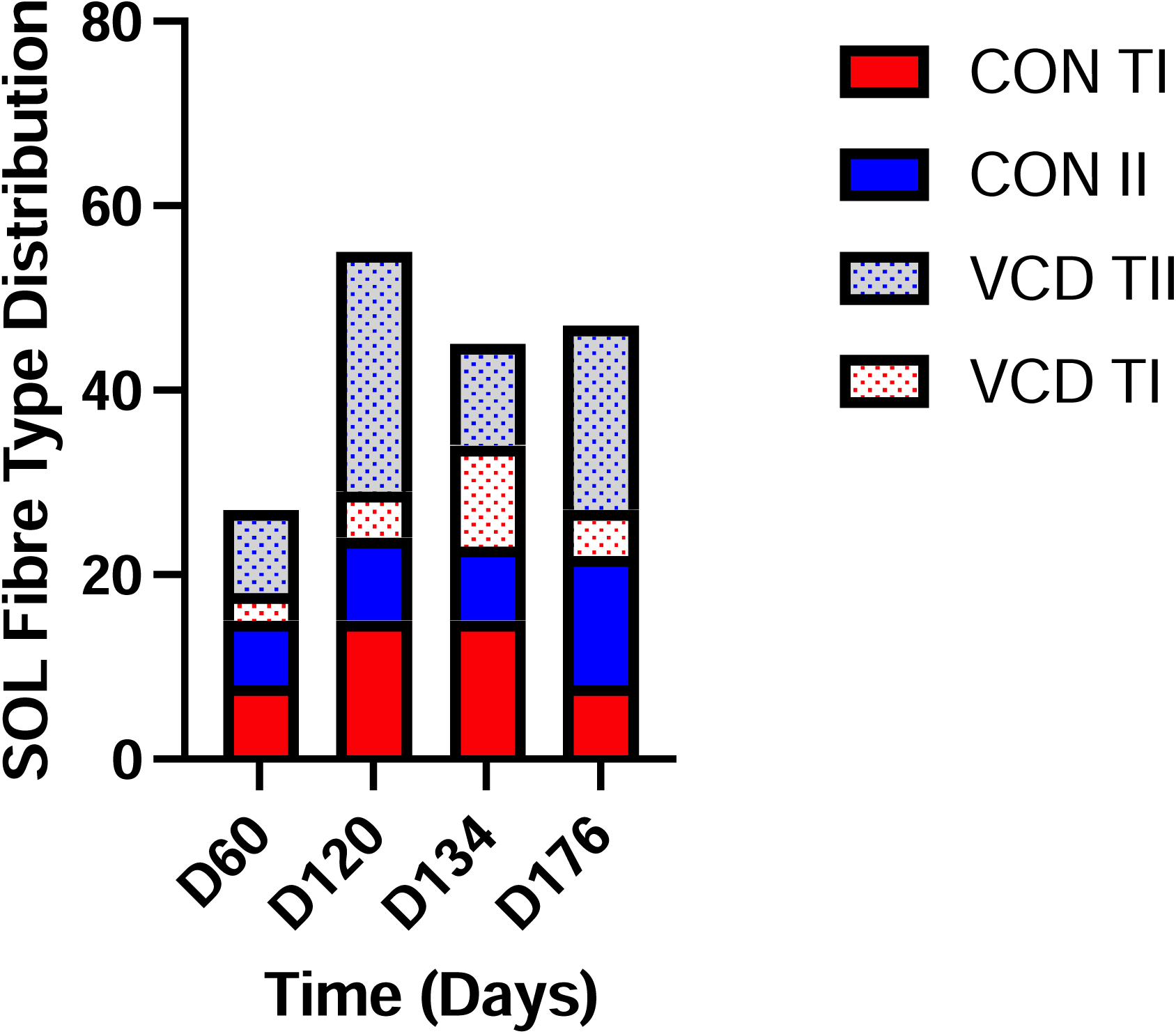
SOL fibre type distribution: The distribution of muscle fibre types within the SOL muscle, illustrating the shift towards more Type II fibres in response to the effects of VCD treatment.

### Extensor Digitorum Longus muscle

#### Absolute force

For absolute force, there was no interaction of group × timepoint (p>0.05) or main effect of group (p>0.05). However, there was a main effect of timepoint (p<0.001), such that fibres in both groups produced ∼36-40% lower absolute force at timepoint D176 compared to all other timepoints (Figure 8A).

**Figure 8.**
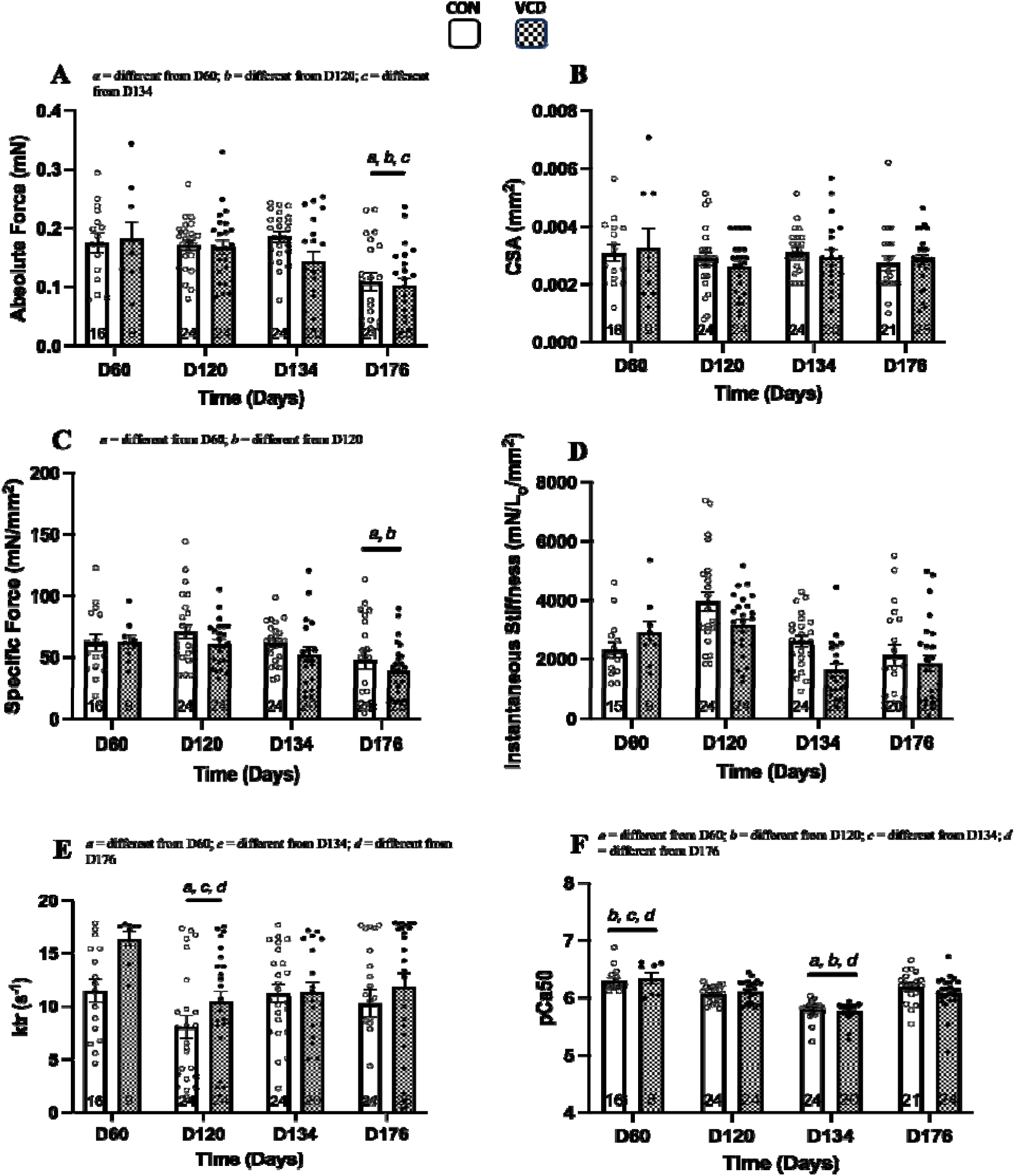
EDL outcome measure results: A-F – Absolute force, CSA, specific force, instantaneous stiffness, redevelopment rate, and pCa_50_ were not different between groups, indicating minimal effects of VCD treatment on muscle contractility on the EDL which highlights the muscle-specific responses to hormonal changes.

#### CSA

For CSA, there was no interaction of group × timepoint (p>0.05), main effect of group (p>0.05) or main effect of timepoint (p>0.05) (Figure 8B).

#### Specific force

For specific force, there was no interaction of group × timepoint (p>0.05) or main effect of group (p>0.05). However, there was a main effect of timepoint (p<0.001), such that fibres at timepoint D176 produced ∼30% and ∼36% lower specific force than at timepoints D60 and D120, respectively (Figure 8C).

#### Instantaneous stiffness

For instantaneous stiffness, there was no interaction of group x timepoint (p>0.05), main effect of group (p>0.05), or main effect of timepoint (p>0.05) (Figure 8D).

#### Ktr

For Ktr, there was no interaction of group × timepoint (p>0.05), or main effect of group (p>0.05). However, there was a main effect of timepoint (p<0.001) such that fibres at timepoint D120 had ∼18-28% slower rate of force redevelopment compared to all other timepoints (Figure 8E).

#### Calcium Sensitivity

For pCa_50_, there was no interaction of group × timepoint (p>0.05), or main effect of group (p>0.05). However, there was a main effect of timepoint (p<0.001) such that fibres at timepoint D60 calcium sensitivity was ∼3-9% higher than all other timepoints and at timepoint D134, calcium sensitivity was ∼5-9% lower than all other timepoints (Figure 8F).

## Discussion

The purpose of the current study was to investigate the effects of VCD-induced ovarian failure in mice on time-course adaptations throughout perimenopause and into late-stage menopause on the contractility of skeletal muscle fibres of the SOL and EDL muscles. Here, we used the VCD model to mimic the gradual ovarian failure similar to menopause in humans and found that for the SOL muscle, force production, fibre size, instantaneous stiffness, and rate of force redevelopment were either improved or were unchanged by late-stage ovarian failure, with fluctuating results at earlier timepoints. These findings indicate that changes to muscle contractility post-onset of ovarian failure do not occur as linearly as expected, and there are likely some underlying compensatory factors maintaining the integrity of muscle mechanical function into late-stage menopause. Moreover, Type I fibres for the SOL improved force capabilities at the onset of- and at late-stage ovarian failure, but interestingly, fibre composition shifted to a greater percentage of Type II fibres from the onset of menopause and at late-stage menopause. For the EDL muscle, we found a uniform decrease in force throughout the menopausal phases and without significant changes in the fibre-type composition of the muscle. Taken together, these data indicate a rather complex relationship between muscle type, fibre type, and ovarian failure, in which a nuanced impact of hormonal changes on muscle physiology and function is observed.

### Alterations to isometric force following ovarian failure are time- and fibre-type dependent

For the SOL muscle across groups, there were no differences in force production capabilities at the perimenopausal phase timepoint (D60). At the onset of- (D120, week ‘0’) and late-stage (D176, week 8) ovarian failure, the VCD groups produced 35% and 29% more force compared to the CON groups, respectively (Figure 1A). At the early-stage (D134, week 2) ovarian failure timepoint, however, the VCD group had lower force production compared to the CON group, which provides critical insight into the temporal effects of ovarian failure, indicating that muscle contractility does not change linearly after ovarian failure (Figure 1A). The D134 timepoint, which is 2 weeks post the onset of ovarian failure, is suggestive of a peak in the negative impact of ovarian failure on muscle function. Using the OVX model, estrogen-deficiency results in a reduction in force production (Cabelka et al., 2019; Cabelka et al., 2021; Greising, Baltgalvis, et al., 2011; Greising, Carey, et al., 2011; Moran et al., 2007; Moran et al., 2006; Peyton et al., 2022) approximately 4-8 weeks following OVX, which differs from our findings at timepoint D176 (8 weeks). However, other studies using the VCD model have found similar results when force was measured at the onset of ovarian failure (D120; week 0) (Mashouri et al., 2024) but not always (Hinks et al., 2024; Hubbard et al., 2023). The initial increase in force observed at the onset of ovarian failure for the VCD group could be attributed, in part, to the relative increase in androgen levels (Kumari et al., 2023; Mashouri et al., 2024), potentially modulated by aromatization, which is the process of converting androgens to estrogens locally in skeletal muscles (Burger, 2001). Additionally, we found that the VCD group had larger CSA compared to the control group at the onset of (D120; week 0) ovarian failure (Figure 2), with no changes for specific force (Figure 3) for that timepoint, indicating that the increase in fibre CSA is likely not due to an increase in myofibrillar content. Our results show that as time progressed from early-stage (D134; week 2) to late-stage (D176; week 8) ovarian failure, the VCD groups maintained force levels while the force levels of the CON groups continuously dropped. Additionally, at early-stage ovarian failure, a reduction in specific force is observed for the VCD group, which is reversed by late-stage ovarian failure, confirming that D134 (week 2) is likely the peak in the negative impact of ovarian failure on skeletal muscle function (Figure 3A). The specific force results at both post-ovarian failure timepoints (D134; week 2 and D176; week 8) can be attributed to changes in the number of bound cross-bridges.

### Rate of force redevelopment (Ktr) did not differ between groups, and calcium sensitivity was higher post-onset of ovarian failure in the SOL

Ktr was not significantly different between groups, which indicates that despite the hormonal changes induced by VCD, the rate of transition of non-force bearing to force bearing cross-bridge states remains largely unaffected throughout the menopausal transition into late state menopause (Figure 5A). Previously we showed an increase in rate of force redevelopment at the onset of ovarian failure (Mashouri et al., 2024). The current study provides a range of time points from pre-ovarian failure to late-stage ovarian failure, and as such, may be better interpreted when considered within the larger framework of the various stages of ovarian failure.

Our results indicate that there was greater calcium sensitivity at all timepoints except for at the onset of ovarian failure (D120; week 0) (Figure 6A), which aligns with previous findings on cardiac muscle (Wattanapermpool, 1998) and SOL muscle (Wattanapermpool & Reiser, 1999) using the OVX model, and the VCD model (Mashouri et al., 2024). The improved calcium sensitivity observed in our study may be due to changes in regulatory protein affinities and post-translational modifications (Gordon & Ridgway, 1993; Gordon et al., 1988), or changes in myosin light chain phosphorylation (Gillis et al., 2016; Lai et al., 2016; Peyton et al., 2022) which require further investigation.

### A shift to an increased proportion of Type II fibres was observed for the SOL muscle, despite the maintenance of force in Type I fibres

The data for the SOL muscle was subdivided by fibre type to tease out mechanisms contributing to changes in force observed at different timepoints. The fibre-typed data revealed that Type I fibres from the SOL maintained or even increased their force output, contrasting with the general decline observed in Type II fibres (Figure 1B). Type I fibres demonstrated resilience or may have compensatory mechanisms that help preserve or enhance their functional capacity despite the hormonal disruptions induced by VCD, suggesting that these fibres might be less susceptible to the adverse effects of ovarian failure, possibly due to differences in metabolism between fibre types, or may be protected because of a greater density of estrogen receptors (ERs). The resilience of Type I fibres to force loss suggests a complex interaction between fibre types and the hormonal environment in skeletal muscles, where at certain stages post-ovarian failure, the adverse effects of ovarian failure might temporarily outweigh the protective or adaptive responses in specific fibre types. Interestingly, a shift towards more Type II fibres was observed in the VCD group (Figure 7), which aligns with previous literature that has reported similar findings using this model (Hubbard et al., 2023; Mashouri et al., 2024) and OVX (Moran et al., 2007), indicating another compensatory adaptation likely to maintain muscle function in response to VCD-induced stress or damage. ERs, particularly ERα are differentially expressed in skeletal muscle and may influence energy metabolism, muscle function, and fibre-type distribution (Cabelka et al., 2019). The interplay between estrogen signalling and ERs could affect muscle function, indicating that differences in ER expression across muscle fibre types may contribute to the observed force disparities of the current study.

### The EDL muscle was seemingly unaffected by VCD-induced ovarian failure

The EDL muscle showed minimal changes in mechanical function. No changes were observed for absolute force, CSA, or specific force (Figure 8A, B, C), indicating that VCD treatment did not significantly alter force production in EDL fibres, except for a decline by D176 and that muscle fibre size remained consistent regardless of VCD treatment. There was a slower rate of force redevelopment at D120 (Figure 8E) while instantaneous stiffness (Figure 8D) remained constant across groups and time, indicating that the number of attached cross-bridges was not significantly affected by VCD treatment. These findings indicate that the EDL muscle may be less sensitive or slower to respond to the hormonal changes compared to the SOL muscle, as previously reported (Greising, Baltgalvis, et al., 2011; Hubbard et al., 2023; Mashouri et al., 2024; Moran et al., 2007).

### Methodological Considerations

The method of fibre typing using SDS-PAGE could introduce selection bias as the "best-looking" fibres are chosen for analysis. This selection might unintentionally favour fibres with more ideal characteristics, potentially skewing the observed distribution of fibre types, as discussed previously (Mashouri et al., 2024). Additionally, the current study includes sexually mature young mice with ovarian failure chemically induced, which may explain some of the differences in results compared to other findings. Age at the onset of ovarian failure, in addition to time from ovarian failure, are key factors that could contribute to impairments in mechanical performance. Strain of rodents, age and time after ovarian failure should be taken into consideration when interpreting and comparing results between studies.

## Conclusion

Muscle contractility across the peri-menopausal transition into late-stage menopause is both muscle and phase-dependent, highlighting the complexity of hormonal impacts on muscle physiology and cellular level mechanical function throughout the lifespan.

## Ethics statement

All procedures were approved by the Animal Care Committee of the University of Guelph (AUP:4714).

## Data accessibility

Supporting data are available upon request.

## Author contributions

Parastoo Mashouri, Jinan Saboune, W. Glen Pyle and Geoffrey A. Power conceived and designed research; Parastoo Mashouri and Jinan Saboune carried out administration of VCD drug; Parastoo Mashouri performed experiments; Parastoo Mashouri analysed data; Parastoo Mashouri, and Geoffrey A. Power interpreted results of experiments; Parastoo Mashouri prepared figures; Parastoo Mashouri and Geoffrey A. Power drafted manuscript; Parastoo Mashouri, Jinane Saboune, W. Glen Pyle and Geoffrey A. Power edited and revised manuscript. All authors have read and approved the final version of this manuscript and agree to be accountable for all aspects of the work in ensuring that questions related to the accuracy or integrity of any part of the work are appropriately investigated and resolved. All persons designated as authors qualify for authorship, and all those who qualify for authorship are listed.

